# The genomic landscape of metastasis in treatment-naïve breast cancer models

**DOI:** 10.1101/777904

**Authors:** Christina Ross, Karol Szczepanek, Maxwell Lee, Howard Yang, Tinghu Qiu, Jack Sanford, Kent Hunter

## Abstract

Metastasis remains the principle cause of mortality for breast cancer and presents a critical challenge because secondary lesions are often refractory to conventional treatments. While specific genetic alterations are tightly linked to primary tumour development and progression, the role of genetic alteration in the metastatic process is not well-understood. To investigate how somatic evolution might contribute to breast cancer metastasis, we performed exome, whole genome, and RNA sequencing of matched metastatic and primary tumours from pre-clinical mouse models of breast cancer. Here we show that in a treatment-naïve setting, recurrent single nucleotide variants and copy number variation, but not gene fusion events, play key metastasis-driving roles in breast cancer. For instance, we identified recurrent mutations in *Kras*, a known driver of tumorigenesis that has not been previously implicated in breast cancer metastasis. The strategy presented here represents a novel framework to identify actionable metastasis-targeted therapies.

## Introduction

Metastatic breast cancer remains the leading cause of cancer-related death among women^1,2^. Of the 1.7 million new cases of breast cancer diagnosed annually worldwide, approximately 30% of patients diagnosed with localized disease eventually present with metastatic lesions in distant organs. While non-metastatic breast cancer has a 5-year survival rate of 99%, metastatic disease reduces 5-year survival to only 25%^2^. Therapeutic strategies to treat localized disease such as molecular profiling and targeted therapy have been increasingly successful, but patients with disseminated disease continue to face much worse outcomes, as metastases are largely insensitive to such treatments^3,4^. Therefore to improve outcome for patients with advanced cancer, specific metastasis-targeted strategies will need to be developed, as will a deeper understanding of the unique biological processes that occur during disease progression^4–6^.

Despite the importance of this process, relatively little is known regarding somatic events that drive the metastatic cascade. The most commonly accepted hypothesis of tumour progression postulates that mutations are acquired over time, resulting in heterogeneous primary tumour tissue composed of distinct subclones^7^. According to this hypothesis, metastatic capability is induced when a subclone acquires all of the necessary secondary genomic alterations to intravasate into the circulation, survive in circulation, arrest and extravasate at a distant site, and colonize that distant organ^7,8^. However, while the primary tumour genetic heterogeneity predicted by this model is widely accepted, evidence of somatic metastasis-driving mutations is lacking^9–11^. Large-scale projects such as The Cancer Genome Atlas (TCGA)^12^ have provided a detailed inventory of oncogenic driver mutations, but no equivalent data set currently exists for metastatic disease. Furthermore, data from our lab and the work of others suggest that metastasis may instead be driven by dynamic epigenetic variation of gene expression programs^13–17^ and there is increasing evidence to favour a model of early metastatic dissemination, which is inconsistent with the somatic evolutionary hypothesis^18–20^. The difficulty of targeting such dynamic gene expression programs and preventing dissemination of cells from clinically undetectable tumours obliges a deeper assessment of the existence of targetable metastasis-driving mutations. In this study we have shown that recurrent single nucleotide variants and copy number variants, but not gene fusion events, occur spontaneously in the absence of therapeutic pressures and drive breast cancer metastasis. Of note, we identified recurrent mutations in the Ras signalling pathway that specifically promoted metastasis *in vivo*.

## Results

### Unique SNVs are enriched in metastases

To complement ongoing human tissue studies and understand the metastatic genomic landscape in a treatment-naïve setting, we performed next generation sequencing on metastatic tissue from pre-clinical mouse models of metastatic breast cancer (Supplementary Fig. 1). In this study we focus on the luminal-like MMTV-PyMT (PyMT) and MMTV-Her2 (Her2) genetically engineered mouse models (GEMMs) of metastatic breast cancer, which model the PI3K activation or the *HER2* amplification seen in 42% or 20% of breast cancer patients, respectively^12,21^. The PyMT model produces palpable primary tumours at a mean latency of 60 days and pulmonary metastases at 100 days in 85% of mice^22^. The Her2 model, produces mammary tumours at a mean latency of 100 days, and pulmonary metastases at 200 days in 60% of mice^23–25^. We crossed the PyMT model (FVB/NJ background) with five mouse strains (FVB/NJ, C57BL/6J, C57BL/10J, CAST/EiJ, MOLF/EiJ) to more closely recapitulate the genetic heterogeneity of human populations. Due to the latency and high variability of the Her2 (FVB/NJ background) model disease progression compared with PyMT, this study only includes Her2 × FVB/NJ data and sequencing of tissues from additional strains is ongoing (Supplementary Fig. 1a).

To investigate whether somatic mutations might contribute to metastatic progression, we performed exome sequencing (exome-seq) of 53 and 12 paired primary tumour (PT) and lung metastases (LM) from the PyMT and Her2 GEMMs, respectively (Supplementary Fig. 1a). Single nucleotide variant (SNV) analysis was performed using three independent SNV-calling algorithms. Genes predicted to harbour SNVs by each algorithm were intersected to identify a common set of potentially mutated genes. We identified 1202 exon SNVs in PT tissue and 1768 in metastatic tissue compared with normal strain-specific gDNA sequence (Supplementary Fig. 2a, b).

Candidate metastasis-driving SNVs were defined by two criteria: 1) enrichment in metastatic tissue but not in primary tumour and 2) presence within the metastatic seeding cell. To identify SNVs likely present in the metastatic seeding cell, only those SNVs found in at least 60% of the metastatic lesion (variant allele frequency between 0.3 to 0.5) were considered as potential metastasis-driver events. This cut-off accounts for the infiltration of non-tumour cells but still requires a heterozygous mutation to be present in at least 60% of the cells within the lesion. The comparison of LM sequences to matched PT identified 196 SNVs in 164 genes common to all 3 SNV calling algorithms (Fig. 1a, Supplementary Fig. 2c & Supplementary Table 1). The 196 genes were then screened to identify recurrently mutated codons. Five genes (*Kras, Shc1, Ccni, Mtch2, Snrk*) were identified with recurrent mutations in the same codon, and again were common to all three algorithms (Supplementary Table 1 and Fig. 1b, d).

**Figure 1.**
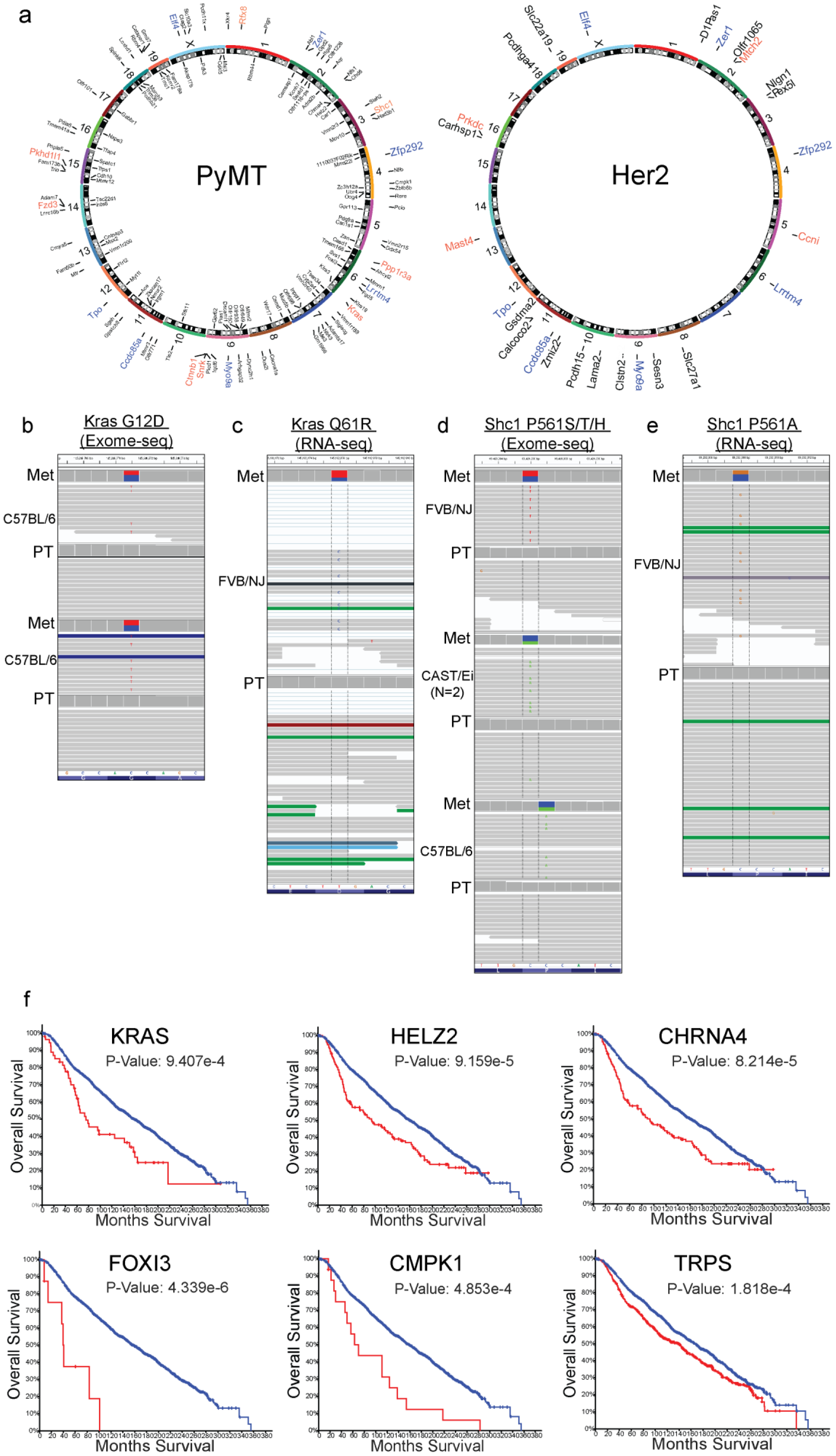
Recurrent metastasis-specific SNVs can be found at high allele frequency. **a.** Circos plots showing names and chromosomal location of genes identified by exome- and RNA-seq with metastasis-specific SNVs (orange: recurrent single strain, blue: recurrent in both strains, and black: singly mutated) from the PyMT model (left) the Her2 model (right). **b-e.** IGV screen shots showing allele frequency in the blue and red boxes, representative SNV reads, mutated codon letter, tissue type, and mouse strain. **b**, Exome-seq identified a C (blue) to T (red) SNV within *Kras* resulting in the G12D amino acid substitution in metastases from two C57BL/6J × PyMT animals. **c**, RNA-seq identified a T (red) to C (blue) SNV within *Kras* resulting in the Q61R amino acid substitution in a metastatic lesion from one FVB/NJ × PyMT animal. **d**, Exome-seq identified C (blue) to T (red) and C (blue) to A (green) SNVs in metastases from three animals (FVB/NJ, CAST/Ei (2 mets from 1 mouse with same SNV), and C57BL/6J × PyMT) resulting in the P561S/T or H amino acid substitutions, respectively. **e**, RNA-seq identified a C (blue) to G (orange) SNV resulting in the P561A amino acid substitution in one FVB/NJ × PyMT animal. **f.** Kaplan-Meier plots generated using METABRIC for the top six genes identified with metastasis-driver SNVs by exome-seq in mice that most significantly stratify patient survival when altered in primary tumour tissue (blue = no CNV, red = CNV present). PT, primary tumour; Met, metastasis.

The SNV calls were then analysed for genes that were recurrently mutated in different codons. A total of 147 genes were identified with mutations observed in only a single animal by all three algorithms. Twelve genes were identified has having been recurrently mutated in different codons in independent animals (Supplementary Table 1). Finally, to prioritize genes for future analysis, RNA sequencing (RNA-seq) data was examined to validate SNVs in tissues analysed by both analyses. An additional instance of the *Shc1* mutated codon was identified, and a potential recurrent mutation in *Rfx8* was found (Supplementary Table 1).

Two genes, *Kras* and *Shc1*, had recurring metastasis-specific mutations in independent animals of the PyMT cohort (Fig. 1b, d). Two PyMT animals carried the oncogenic activating *Kras* G12D mutations (Fig. 1b), and an additional three PyMT animals carried different nucleotide substitutions all within the proline 561 or 451 codon (isoform a or b respectively) of *Shc1*, resulting in P561S, P561T, or P561H (Fig. 1d). For those samples with gDNA or RNA still available (4 out of 7), Sanger sequencing validated the presence of *Kras* and *Shc1* SNVs in the original metastatic tissues analysed by exome- and RNA-seq, and in some cases in additional metastases from the same animal (Supplementary Table 2). The presence of recurrent metastasis-enriched SNVs in several animals in the absence of therapeutic pressures suggests that specific coding mutations may play a fundamental role in driving metastasis.

To expand the search for additional genes with recurrent metastasis-enriched mutations, we screened RNA-seq data from a cohort of 42 matched primary tumours and metastatic pairs from 40 PyMT and 2 Her2 animals (Supplementary Figure 1a). The RNA-seq data included 17 PyMT animals also analysed by exome-seq plus an additional 25 independent animals (2 Her2 and 23 PyMT) (Supplementary Figure 1a). First, the 164 genes with high-probability SNVs identified by exome-seq were screened against the RNA-seq data to validate exome-seq SNV predictions and identify additional animals with mutations in these genes (Supplementary Table 1 & 2). For animals used in both analyses, the predicted SNVs identified by exome-seq were validated by the RNA-seq data given the gene was transcribed, suggesting that the exome-seq filtering criteria correctly identified bona fide SNVs. One additional metastasis-enriched *Kras* mutation (Q61R; Fig. 1c) and an additional mutation in *Shc1* P561 (P561A; Fig. 1e) were observed in two animals from the RNA-seq-only cohort. Unexpectedly, in addition to a single metastasis-enriched *Ctnnb1* oncogenically-activating SNV (S45P)^26^ observed in exome-seq data, RNA-seq data revealed three more independent animals possessing metastasis-specific *Ctnnb1* S45F SNVs and another two with *Ctnnb1* point mutations that are also often observed in human tumours (K335N, N387K) (Supplementary Table 1). Sanger sequencing validated the presence of the SNVs for *Ctnnb1* exome-seq S45P and RNA-seq N387K mutations (Supplementary Table 2). Insufficient sample material prevented Sanger validation of the remaining RNA-seq-identified SNVs. Overall, for samples with available material, the rate of validation by Sanger sequencing for metastasis-specific SNVs was approximately 80% (8 out of 10) (Supplementary Table 2 and Supplementary Fig. 1). Finally, Sanger sequencing of matched pairs of tumours and metastases from an additional 22 animals not used for RNA- or exome-seq was used to survey for 11 of the SNVs identified by exome- and RNA-seq. This analysis identified yet another *Shc1* P561H mutation, but no additional samples containing the other 10 mutations were found.

### Copy number variation drives metastasis and reduces overall survival among human breast cancer patients

Using the METABRIC sequencing dataset of human breast cancer primary tumours, we assessed the relevance to human disease of genes possessing metastasis-enriched SNVs identified in the PyMT and Her2 mouse models. This analysis revealed that many genes possessing SNVs in our preclinical mouse models were amplified in human tissue, and 18% (23 out of 123 human orthologs) of such alterations were significantly associated with poor patient outcome (Fig. 1f and Supplementary Fig. 4 & 5).

Since human primary tumours possessed gene amplification or deletion in addition to point mutations, we analysed the PyMT primary tumour/metastasis exome-seq pairs for potential copy number variations (CNVs) that may contribute to metastatic spread. To determine allele-specific gain or loss we used only those samples generated from progeny of the FVB/NJ-PyMT outcross as their genomes contained an FVB and non-FVB (C57BL/6J, C57BL/10J, CAST/EiJ, or MOLF/EiJ) allele that could be readily differentiated. Therefore, pure FVB/NJ-PyMT and FVB/NJ-Her2 tumour/metastasis pairs were not analysed due to the lower confidence of CNV calling on a homozygous genetic background. Based on these criteria, we performed CNV analysis using exome-seq data from 30 of the primary tumour/metastasis pairs in our cohort. We first identified CNV in primary tumours and metastases compared to normal, and then analysed this data for metastasis-specific CNVs (Supplementary Tables 3-7). Amplification and deletion events were primarily restricted to the F1 animals of crosses with C57BL/6J and MOLF/EiJ, with recurrent metastasis-specific CNVs on chromosomes 2, 4, 9, and 10 (Fig. 2a). Examination of the putative CNVs suggested that the C57BL/6J alleles were specifically lost in the F1 hybrids, while FVB/NJ alleles were under-represented in most of the CNVs in the MOLF/EiJ F1 hybrids. In addition, the chromosome 4 and 9 CNVs overlap with regions of the genome we have previously demonstrated to harbour inherited metastasis susceptibility genes^27–29^, suggesting these genomic intervals may contain important metastasis-associated factors. The Genomic Regions Enrichment of Annotations Tool (GREAT)^30^ was used to identify the genes associated with recurrent metastasis-specific CNVs. 371 genes were associated with recurrent regions of CNV in PyMT metastatic lesions, and 46 of these genes possessed CNVs in two or more outcross strains (Supplementary Table 8 and Supplementary Fig. 4). The METABRIC dataset was then screened again to assess if CNVs within any of the 46 genes were significantly associated with breast cancer patient survival. The amplification of approximately 17% (8 out of 46) of the genes queried was significantly associated with worse patient outcome, including *EBAG9* and *PKHD1L1*, located on mouse chromosome 15 and *PEX2*, and *ZFHX4*, located on mouse chromosome 2 (Fig. 2b), which are all located on human chromosome 8 between q21.13 and q23.2. This analysis reveals that in addition to metastasis driver SNVs, metastatic cells may also acquire specific CNVs to support spread to a secondary site.

**Figure 2.**
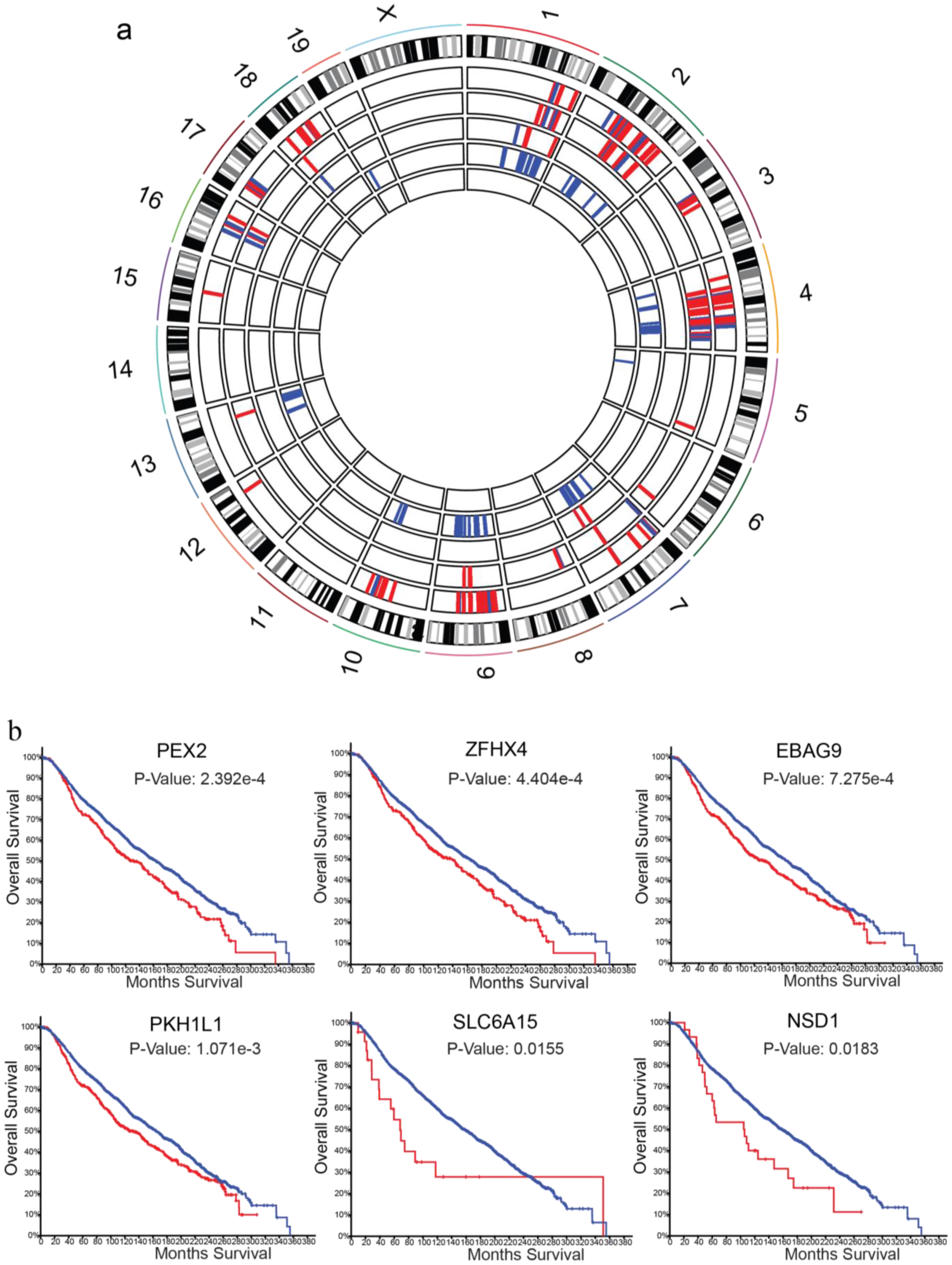
Metastasis-specific CNVs and SNVs in the PyMT and Her2 mouse models. **a.** Circos plot representing metastasis-specific genomic alterations. The outermost track shows the mouse chromosomes and established G-banding. The remaining five inner tracks show deleted (and amplified for MOLF/EiJ only) genomic regions in F1 PyMT mice resulting in enrichment of the non-FVB/NJ allele (red) or enrichment of FVB/NJ allele (blue). From the outermost track the strain order is as follows: MOLF/EiJ amplification, MOLF/EiJ deletion, CAST/EiJ deletion, C57BL/6J deletion, C57BL/10J deletion. **b.** Kaplan-Meier plots generated using METABRIC for six out of eight genes identified as metastasis drivers by exome-seq CNV analysis in mice that significantly stratify patient survival when altered in primary tumour tissue and reside on human chromosome 8 (blue = no CNV, red = CNV present).

### Fusion genes are not enriched in metastases

In a large-scale analysis of 560 human breast cancer samples, Nik-Zainal et al. discovered frequent variation in genomic structure in primary tumour tissue^31^. To identify any potential targetable gene fusion events that may contribute to metastatic progression, we analysed the RNA-seq data (Supplementary Fig. 1) using the deFuse algorithm to identify discordant paired end alignments and thus putative gene fusion events specific to metastatic tissue^32^. To reduce false positive results, putative alternative splice events from adjacent genes were excluded from the analysis to eliminate rare transcriptional read-through products. Forty-four putative fusion transcripts were detected in the primary tumours by this analysis. Seventy-one fusion transcripts were detected in the metastases, with 17 fusions in common with the primary tumours and 54 unique to the metastatic lesions. The 54 metastasis-specific putative fusion events involved a total of 85 genes. Sixteen genes were associated with putative fusion events in more than one animal or involved genes with multiple fusion partners (Fig. 3a). However, manual curation of the putative recurrent fusion events revealed that 7 of the 15 breakpoints occurred in regions of repetitive elements within introns or UTRs, suggesting potential alignment artefacts. Moreover, for the highly expressed genes (*Csn3, Trf*), the variant transcripts were represented at <1% of the total read count, suggesting that these transcripts were either not required for maintenance of the metastatic lesions, rare aberrant transcripts, or were artefacts of the RNA-seq/deFuse analysis.

**Figure 3.**
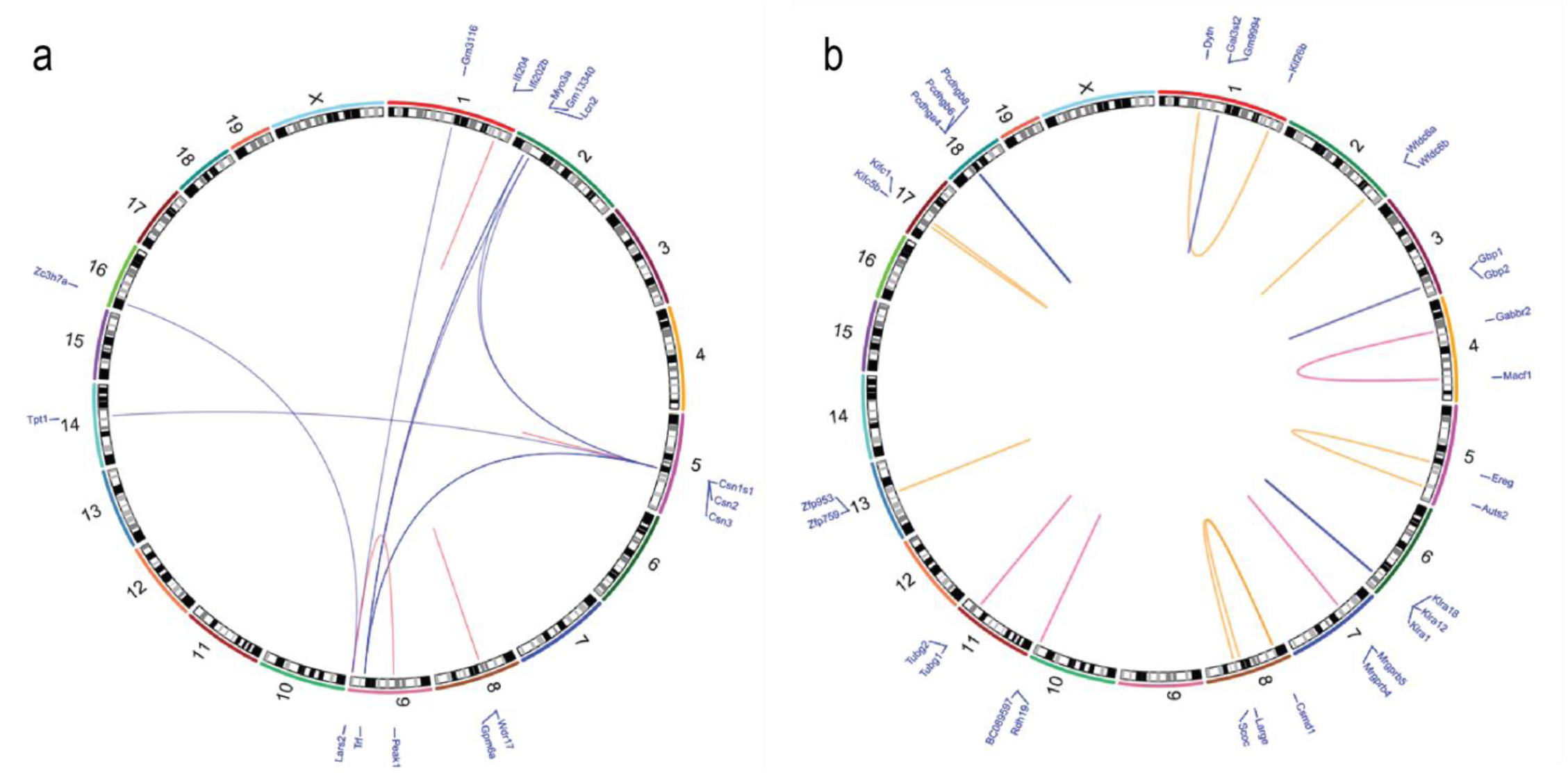
Metastasis-specific structural variants do not drive metastasis in pre-clinical breast cancer mouse models. Circos plots representing deFuse2 analysis of RNA-seq data for the 41 matched primary and metastatic PyMT tumours. BreakDancer algorithm analysis of 10x WGS data from 20 matched pairs of primary and metastatic lesions from FVB/NJ PyMT × MOLF/EiJ and FVB/NJ PyMT × CAST/EiJ animals. For both plots, the names of the genes associated with predicted structural variations are listed in blue in the outermost track by their corresponding chromosome numbers, which are found in the second track. The inner loops represent breakpoints within repetitive elements at both sites (pink loops), repetitive element at one site (tan loops), or within single copy sequence at both sites (blue loops).

Additionally, we performed 10x whole genome sequencing (WGS) on 20 matched pairs of primary and metastatic lesions from FVB/NJ-PyMT×MOLF/EiJ and FVB/NJ-PyMT × CAST/EiJ animals. This data set was comprised of tissue from 16 of the animals represented in the exome-seq dataset and 4 additional animals (Supplementary Fig. 1a). MOLF/EiJ and CAST/EiJ F1 animals were used for this analysis as their genomes are significantly polymorphic compared to FVB/NJ, allowing for allele-specific identification. SNVs predicted by exome-seq were confirmed in all of those animals also included in the WGS analysis (Supplementary Table 2). Putative coding-related structural variants such as insertion-deletions (indels), inversions, and translocations were identified using the BreakDancer algorithm^33^ by limiting the analysis to within transcripts and requiring a minimum of three variant reads per sample (Supplementary Table 9). The resulting gene fusions were found to be predominantly intrachromosomal and tended to cluster within regions of gene families, suggesting potential alignment artefacts. To reduce this potential artefact, a threshold of a minimum of 10 kb between putative fusion partners was introduced, resulting in 18 putative fusion events involving 31 genes observed in either more than one animal or with multiple fusion partners (Fig. 3b). No overlap between the putative structural variants observed from the deFuse and BreakDancer analysis was observed. All structural variations predicted by BreakDancer under these conditions were intrachromosomal, and 11 of the 18 predicted recurrent structural variants occurred between 2 members of the same gene family. Of the remaining seven, two of the remaining predicted structural variation breakpoints were within repetitive elements at both sites (Fig. 3b, pink loops). Two additional predicted structural variations contained a repetitive element at one of the breakpoints (Fig. 3b, tan loops), while three were within single copy sequence at both sites (Fig. 3b, blue loops). These results suggest that structural variation resulting in fusion transcripts does not play a significant role in the metastatic progression of the MMTV-PyMT mammary tumour model.

### Single nucleotide variants drive metastasis

The presence of recurrent oncogenic mutations in *Kras* and *Ctnnb1*, in addition to the recurrent mutations of the codon of *Shc1*, are consistent with a potential metastasis-driving function of these genes in the PyMT mouse model. Interestingly, previous analysis of a panel of metastatic mouse mammary tumour cell lines revealed recurrent mutation of *Kras* in several cell lines^34^. Moreover, SNVs within *KRAS* in the primary tumours of the METABRIC data set, although rare (0.6% of patients), was significantly associated with poor survival in human breast cancer (Fig. 4a, b). Together, these observations suggest that this signalling axis may play an important role not only in tumorigenesis, but also in metastatic progression of human malignancies and therefore warrants further investigation. *CTNNB1* and *SHC1* were amplified (0.3% and 22% of patients, respectively), but were not included in the targeted sequencing performed on 173 cancer-related genes in METABRIC. Therefore, the mutational burden for these genes in patient primary tumours within this data set is unknown. We thus focused on *Kras* as a potential breast cancer metastasis driver gene. Wildtype (WT) or *Kras* G12D constructs were expressed in two independent *Kras* WT mouse mammary tumour cell lines: 4T1, which is derived from a spontaneous BALB/c mammary tumour, and MET1, derived from the MMTV-PyMT model (Supplementary Fig. 7a)^34^. *Kras* WT-, *Kras* G12D-, or control empty vector (EV)-transduced cells were implanted orthotopically in syngeneic mice and primary tumour weight and the number of pulmonary metastases were assessed after four weeks. Expression of *Kras* WT and G12D increased tumour burden compared to EV in the MET1 line and a small but non-significant increase in tumour growth was also observed with the 4T1 line (Fig. 4c, f). Therefore, to assess metastatic capacity specifically, the number of metastatic nodules (Fig. 4d, g) was normalized to primary tumour weight for each mouse. This analysis revealed a significant increase in metastatic capacity of *Kras* G12D-expressing cells compared to both *Kras* WT and EV cells in both lines (Fig. 4e, h). To further test how modulation of *Kras* G12D affects metastatic potential, we utilized the 6DT1 mouse mammary cancer cell line that bears an endogenous homozygous *Kras G12D* mutation^34^. Despite the short-lived effects of siRNA, knock down of *Kras* in 6DT1 cells reduced metastasis to the lungs in spontaneous metastasis assays compared to a non-targeting siRNA control (Fig. 4i-k and Supplementary Fig. 7b). This result was significant before controlling for primary tumour weight, and trending after normalization (p=0.1) (Fig. 4j, k). These results are consistent with a function for constitutively activated *Kras* as a metastasis driver gene in breast cancer.

**Figure 4.**
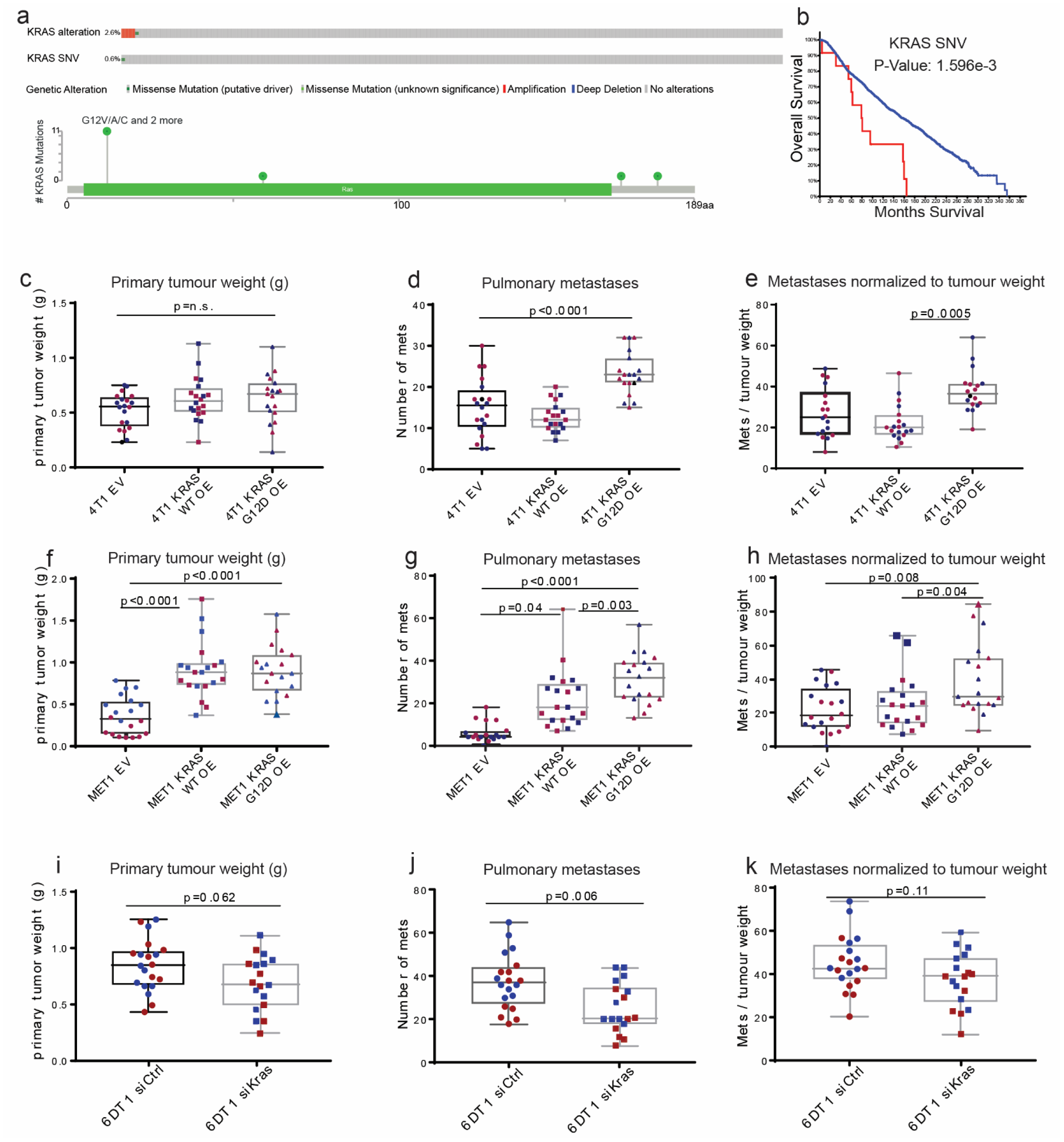
The *Kras* G12D mutation drives breast cancer metastasis. **a.** METABRIC sample data showing that *KRAS* is altered in 2.7% and mutated in 0.6% of primary tumours from breast cancer patients. **b.** Kaplan-Meier analysis of patients with SNVs (red) in *KRAS* vs wildtype (blue). **c-k.** Primary tumour weights, numbers of metastases, and metastases normalized to primary tumour weight from orthotopic injection of 4T1 cells transduced with empty vector, *Kras* wildtype, or *Kras* G12D (**c-e**), MET1 cells transduced with empty vector, *Kras* wildtype, or *Kras* G12D (**f-h**), or 6DT1 cells transfected with siControl or si*Kras* (**i-k**). Each assay was performed twice with 10 mice per group (red and blue points), box whiskers represent min and max points, box boundaries represent the 25th to 75th percentiles, and the horizontal line within the box represents the median. p<0.05 = significant.

## Discussion

Breast cancer metastasis is the primary cause of breast cancer related mortality, and due to the disparate biology of metastatic lesions compared to the original tumour there are few options for clinical intervention ^4,5,35–38^. In this study we hypothesized that somatic evolution of the tumour cell genome may drive metastasis and also may provide novel therapeutic targets. However, to date a robust sequencing dataset of paired treatment-naïve primary and metastatic tumour samples that is large enough to identify recurrent mutations genomic events is not available for such analysis. A key obstacle in this progress is the difficulty of obtaining appropriate tissue samples. Metastatic lesions are usually not surgically removed, and data obtained from resected lesions are usually confounded by the application of neo-adjuvant and adjuvant therapies, which may select for events associated with treatment resistance rather than metastasis^39^. Therefore, in this study we have used mouse models to overcome these limitations and as a hypothesis generation and testing platform for subsequent validation and characterization in human experimental systems.

Here we have performed exome, whole genome, and RNA sequencing on a total of 94 pairs of treatment-naïve primary and metastatic tumours^21,40^ of the luminal-like PyMT and Her2 genetically engineered mouse models. Our analyses identified multiple recurring metastasis-specific SNVs in lesions at high allele frequency in a treatment-naïve setting, indicating that the variants were likely present in the metastatic seeding cell and drove metastasis from a subclone within the primary tumour. Indeed, we show here that the recurrent *Kras* G12D mutation observed in metastatic lesion from 2 PyMT mice increases the metastatic efficiency of mouse mammary cancer cell lines in an orthotopic implantation model. Furthermore, by overlapping sequencing analysis of several tissues, 96% (47 out of 49) of SNVs identified by Exome-seq were confirmed in an orthogonal method (Supplementary Table 2). Taken together our data support the Nowell hypothesis of tumour evolution as a driver of metastasis^7^.

Interestingly, unlike oncogenic driver mutations, we did not observe a high percentage of lesions with mutations in any specific gene, suggesting that there may be many metastasis drivers, each contributing a small fraction of the overall metastasis-driving function within a population. We did, however, observe several SNVs located in genes functioning within the Ras signalling pathway. These SNVs were observed in ∼15% (14 (3-*Kras*, 5-*Shc1*, 6-*Ctnnb1*) out of 94) of the animals in this study, which implicates the *Kras* pathway as an important metastasis signalling axis in this model. The KRAS G12D mutation is a well-studied oncogenic driver leading to the development of aggressive lung and colorectal cancers^41–43^. However, KRAS is not considered a driver of human breast cancer development and therefore its metastasis-specific recurrence in this model was unexpected ^44^. Additionally, the exact role of the *Shc1* P561X mutation is unknown, but the localization of this mutation to the SH2 domain of SHC1 is consistent with activating function and warrants further investigation ^45^.

The SNV data reported here also suggest that there may be many ways within metastasis-associated pathways to activate a pro-metastatic function. The identification of *Kras* and *Ctnnb1* as recurrently mutated metastasis drivers was unanticipated due to the lack of association observed in human breast cancer. The presence of known, well-studied activating mutations in these genes within metastasis-founding cells, however, strongly implicates the Kras-Ctnnb1 signalling axis as a critical component of metastatic progression, at least in the PyMT model system. The activation of the *Ctnnb1* pathway has been previously linked to breast cancer progression and thus our data corroborates the importance of this network^46^, and suggests an additional therapeutic opportunity to intervene in metastasis establishment and progression.

We have also identified recurrent metastasis-specific regions of CNV that may contribute metastatic spread. Several studies have shown divergent CNV between primary and metastatic tissue from small patient cohorts^47–50^, and through our use of the treatment-naive PyMT animal model we have further shown that these genomic alterations occur through natural tumour progression and can drive metastasis. Additionally, our data identified several genes associated with metastasis-specific CNV that, while distributed across the mouse genome, are all localized within human chromosome 8q. Amplifications on chromosome 8q are associated with tumour progression and worse outcomes in many cancer types, suggesting that our analysis appropriately captures genetic alterations observed in human patients^51–56^. This association suggests that our strategy for the identification of metastasis-driving CNVs aligns with human biology, revealing a robust tool for the identification of clinically-relevant metastatic drivers. We anticipate that with additional pairs, strain-specific and driver-specific metastasis-driver CNVs will be distinguished.

In this study we compare the metastasis-specific genomic alterations from mouse models to human sequencing data produced by the METBRIC study. Despite the large number of patients included in this study, it is somewhat limited as it largely reports survival associated primary tumour CNVs, and only 173 genes underwent targeted sequencing^57,58^. As we continue to sequence subsequent mouse models of metastatic breast cancer such as Her2, MMTV-Myc, and C3-Tag we anticipate additional human datasets will become available for expanded mouse-to-human SNV comparisons.

In summary, by performing a survey of the genomic landscape of metastatic progression in two preclinical mouse models, we have identified recurrent, clinically-relevant mutations in genes and pathways that drive metastasis. Metastasis is the leading cause of cancer-related death in breast cancer patients, with over 30% of breast cancer patients progressing to metastatic disease which currently averages just 25% 5-year survival. By using the strategy outlined here, the continued characterization of preclinical mouse models of metastasis will provide further insight into the associations between clinical subtypes, primary tumour drivers, and metastasis-driving genomic alterations, ultimately creating new opportunity for the generation of metastasis-targeted therapies.

## Methods

### Ethics statement

The research described in this study was performed under the Animal Study Protocol LCBG-004 and LPG-002, approved by the National Cancer Institute (NCI) Animal Use and Care Committee. Animal euthanasia was performed by cervical dislocation after anaesthesia by Avertin.

### Genetically engineered mouse models

FVB/N-Tg(MMTV-PyVT)634Mul/J (PyMT) and FVB/N-Tg(MMTVneu)202Mul/J (Her2) male mice were obtained from Jackson Labs. Male PyMT mice were crossed with female wild type FVB/NJ, MOLF/EiJ, CAST/EiJ, C57BL/6J, and C57BL10/J mice also obtained from Jackson Labs. Male Her2 mice were crossed with female wild type FVB/NJ mice. All female F1 progeny were genotyped by the Laboratory of Cancer Biology and Genetics genotype core for the PyMT or Her2 gene and grown until humane endpoint. Mice were euthanized using intraperitoneal Avertin to anesthetize followed by cervical dislocation. Primary tumour, metastatic nodules, and normal (tail) tissue were isolated immediately following euthanasia and snap frozen in liquid nitrogen. Tissue samples were then stored at -80°C.

### gDNA isolation

Primary tumour tissue was ground on dry ice and small fragments were taken for gDNA isolation. Whole metastases were used for gDNA isolation. Tissue was lysed using Tail Lysis Buffer (100 mM Tris-HCl pH 8.0, 5 mM EDTA, 0.2% SDS, 200 mM NaCl, 0.4 mg/ml proteinase K) at 55°C overnight. Samples were then placed in a shaking (1400 rpm) heat block for 1 hour at 55°C. RNaseA (Thermo Fisher Scientific) was added (2 mg/ml final) and lysates were incubated on the bench for 2 minutes. gDNA was then isolated using the ZR-Duet DNA/RNA MiniPrep kit (Zymo Research).

### Sequencing and analysis

All analyses were carried out on the NIH Biowulf2 high performance computing environment. All analyses were performed using software default parameters if not otherwise specified.

### Exome sequencing

Exome sequencing was performed by the NCI Center for Cancer Research (CCR) Genomics Core and the NCI Illumina Sequencing Facility. Exome libraries were prepared using an Agilent SureSelect^XT^ Mouse All Exon Kit for target enrichment. Libraries were barcoded and pooled before sequencing on an Illumina HiSeq3000 or HiSeq4000 to an average depth of 40x. Samples were trimmed of adapters using Trimmomatic software (0.39). The trimmed reads were aligned to the mm10 reference mouse genome using BWA (0.7.17) or Bowtie (2-2.3.5.1) mapping software. SAMtools (1.9) mpileup, GATK (3.8-1) Mutect2, and Strelka (2.7.1) were used to identify potential variants and the variants were filtered for known polymorphisms from mgp.v5 (ftp://ftp-mouse.sanger.ac.uk/current_snps/mgp.v5.merged.snps_all.dbSNP142.vcf.gz) and variants with Phred-scaled quality score of < 30 were removed. Annotation was performed by Annovar (2018-04-16). High probability SNVs were identified after reads were visualized in IGV and false positives were removed from the data. Select SNVs validated by Sanger sequencing performed by the CCR Genomics Core. Copy number variation was performed for the F1 tumours using the R (3.6.0) packages BubbleTree (2.15.0) and cn.mops (1.31.0), followed by filtering for metastasis-specific events and manual curation. We used BubbleTree to do allele-specific and strain-specific copy number analysis, which required genotype data, and cn.mops for general copy number analysis, which doesn’t require F1 cross experiment data. The recurrent CNVs were defined as CNVs present in multiple samples.

### Whole genome sequencing

Library preparation was performed using the Illumina TruSeq DNA Sample Prep Kit FC-121-1001. Samples were barcoded, pooled, and sequenced on an Illumina HiSeq4000 to a depth of ∼10x per sample. Samples were trimmed of adapters using Trimmomatic (0.39) software before the alignment. The trimmed reads were aligned with the mm10 reference mouse genome mm10 using BWA alignment. SNV calls were performed as described for exome sequencing. Structural variation analysis was performed using BreakDancer (1.4.5), followed by filtering for metastasis-specific events and manual curation. For breakpoints located within or near a gene, we filtered structural variations (SVs) by limiting the breakpoint within 10 kb of a gene. CNV analysis was performed using cn.mops as described for exome sequencing.

### RNA sequencing

RNA fusion transcripts were identified using deFuse (0.8.1) software. We used the filtered fusion transcripts. The putative fusion transcripts were further analysed by comparing with WGS BreakDancer data to identify fusion transcripts that were supported by genomic changes. Mutant alleles from RNA-seq data were identified using SAMtools (1.9). The sequence reads in pileup format were generated using the SAMtools mpileup utility for BAM file spanning the SNVs. We extracted the reference allele and mutant allele counts for each sample using a custom-built Perl (v5.24.3) script and R (3.6.0) script.

### Sanger sequencing

Polymerase chain reaction (PCR) was performed using AmpliTaq Gold polymerase (Thermo Fisher Scientific) according to the manufacturer’s instruction. PCR products were then purified by gel electrophoresis using the QIAquick Gel Extraction Kit (Qiagen). The Sanger reaction and electrophoresis for the forward and reverse sequences of each purified amplicon was performed by the CCR Genomics Core. Sequences were manually assessed for single nucleotide variations using Geneious software. If the original sample was RNA, then cDNA was synthesized by reverse transcription using the iScript cDNA Synthesis Kit (Bio-Rad).

### Genomic Regions Enrichment of Annotations Tool (GREAT) analysis

BED files containing the recurrent regions of genomic gain or loss for each mouse non-FVB mouse strain were loaded into the GREAT tool website using the default settings for gene assignment. GREAT calculates statistics by associating genomic regions with nearby genes and applying the gene annotations to the regions.

### Cell culture

Mouse mammary carcinoma cell lines 6DT1, 4T1, and MET1 were a generous gift from Dr Lalage Wakefield (NCI, Bethesda, MD). All cell lines were cultured in Dulbecco’s Modified Eagle Medium (DMEM), supplemented with 10% Fetal Bovine Serum (FBS), 1% Penicillin and Streptomycin (P/S) (complete DMEM), and 1% L-Glutamine (Gibco), and maintained at 37°C with 5% CO2. Short interfering RNA (siRNA)-mediated knockdown and overexpression cells were cultured in the same conditions with an addition of 10 μg/ml puromycin and 5 μg/ml blasticidin, respectively.

### Plasmid constructs

Lentiviral pDEST Gateway Entry clones expressing *Kras* WT or *Kras* G12D with a C-terminal myc tag, and under the control of the *Pol2* promoter were obtained from the NCI Ras Initiative. An empty lentiviral pDEST Gateway Entry clone (Thermo Fisher Scientific) was used as the empty vector control.

### Virus transduction

1 × 10^6^ 293T cells were plated in 6 cm dishes 24 hours prior to transfection in P/S-free 10% FBS DMEM media. Cells were transfected with 1 μg *Kras* or control expression plasmid and 1 μg of viral packaging plasmids (250 ng pMD2.G and 750 ng psPAX2) using 6 μl of X-tremeGENE 9 transfection reagent (Roche). After 24 hours of transfection, media was refreshed with complete DMEM. The following day, virus-containing supernatant was passed through a 45-μm filter to obtain viral particles, which were then transferred to 100,000 4T1 or MET1 cells. The viral media was removed and fresh complete DMEM was added 24 hours post-transduction. Finally, the cells were selected with 10 μg/ml puromycin- or 5 μg/ml blasticidin-containing complete DMEM beginning 48 hours after transduction.

### siRNA transfection

6DT1 cells were plated in in P/S-free 10% FBS DMEM media. 24 hours after plating, cells were transfected with AllStars Mouse Negative Control siRNA (Qiagen) or si*Kras* (si*KRAS*_234 as described by Yuan et al.^59^) using RNAiMAX (Invitrogen).

### *In vivo* metastasis experiments

Female virgin FVB/NJ or BALB/cJ mice were obtained from Jackson Laboratory, and athymic NCI Nu/Nu from NCI Frederick at 6–8 weeks of age. Two days prior to *in vivo* experiments, cells were plated at 1 × 10^6^ cells/condition into T-75 flasks (Corning) in non-selective DMEM. A total of 100,000 cells per mouse was injected into the fourth mammary fat pad of FVB/NJ (MET1 and 6DT1 cells), BALB/cJ (4T1 cells) mice. The mice were euthanized between 28–30 days post-injection. Primary tumours were resected, weighed, and lung metastases counted. Statistical significance was calculated with a Kruskal-Wallis test followed by Conover Inman test. All animal experiments were performed in compliance with the National Cancer Institute’s Animal Care and Use Committee guidelines.

### Western blotting

Protein lysate from 1 × 10^6^ cells were extracted on ice using Golden Lysis Buffer (10 mM Tris pH 8.0, 400 mM NaCl, 1% Triton X-100, 10% Glycerol+Complete protease inhibitor cocktail (Roche), phosphatase inhibitor (Sigma)). Protein concentration was measured using a BCA Protein Assay Kit (Pierce) and analysed on the Versamax spectrophotometer at a wavelength of 560 nm. Appropriate volumes containing 20 μg of protein lysates combined with NuPage LDS Sample Buffer and NuPage Reducing Agent (Invitrogen) were run on 4–12% (or otherwise indicated) NuPage Bis-Tris gels in MOPS buffer. Proteins were transferred onto a PVDF membrane (Millipore), blocked in 5% milk (TBST + dry milk) for 1 hour and incubated in the primary antibody (in 5% milk) overnight at 4°C. Membranes were washed with 0.05% TBST (TBS + 5% Tween) and secondary antibody incubations were done at room temperature for 1 hour. Proteins were visualized using Amersham ECL Prime Western Blotting Detection System and Amersham Hyperfilm ECL (GE Healthcare).

The following primary antibodies were used: mouse anti-Actin (1:10,000; Abcam), mouse anti-myc-tag (1:1000; Cell Signaling Technology), mouse anti-KRAS (1:1,000; Sigma). Secondary antibodies, including goat anti-rabbit (Santa Cruz) and goat-anti-mouse (GE Healthcare), were used at a concentration of 1:10,000.

### Statistics

Statistical significance between groups in *in vivo* assays was determined using the Mann-Whitney unpaired nonparametric test using Prism (version 5.03, GraphPad Software, La Jolla, CA). Statistical significance between samples in RT-qPCR analysis was determined by an unpaired t test, also using Prism.

## Data Availability Statement

All sequence data that supports the findings of this study will be deposited in a public repository and the accession codes will be made available prior to publication.

## Acknowledgements

This work utilized the computational resources of the NIH HPC Biowulf cluster. (http://hpc.nih.gov).

This research was supported by the Intramural Research Program of the NIH, National Cancer Institute, Center for Cancer Research.

## Author Contributions

Christina Ross, Karol Szczepanek, and Kent Hunter conceived of the presented idea. Christina Ross and Karol Szczepanek set up GEMM breedings, tissue isolation, and gDNA / RNA isolation. Christina Ross also performed tissue culture, *in vivo* metastasis assay, and manuscript preparation. Maxwell Lee and Howard Yang performed bioinformatics and statistical analysis. Tinghu Qiu performed PCR and Sanger sequencing, Jack Sanford performed gDNA and RNA isolation from GEMM tissue isolates. Kent Hunter provided resources and intellectual contribution.

## Supplementary Figures

**Supplemental Figure 1.**
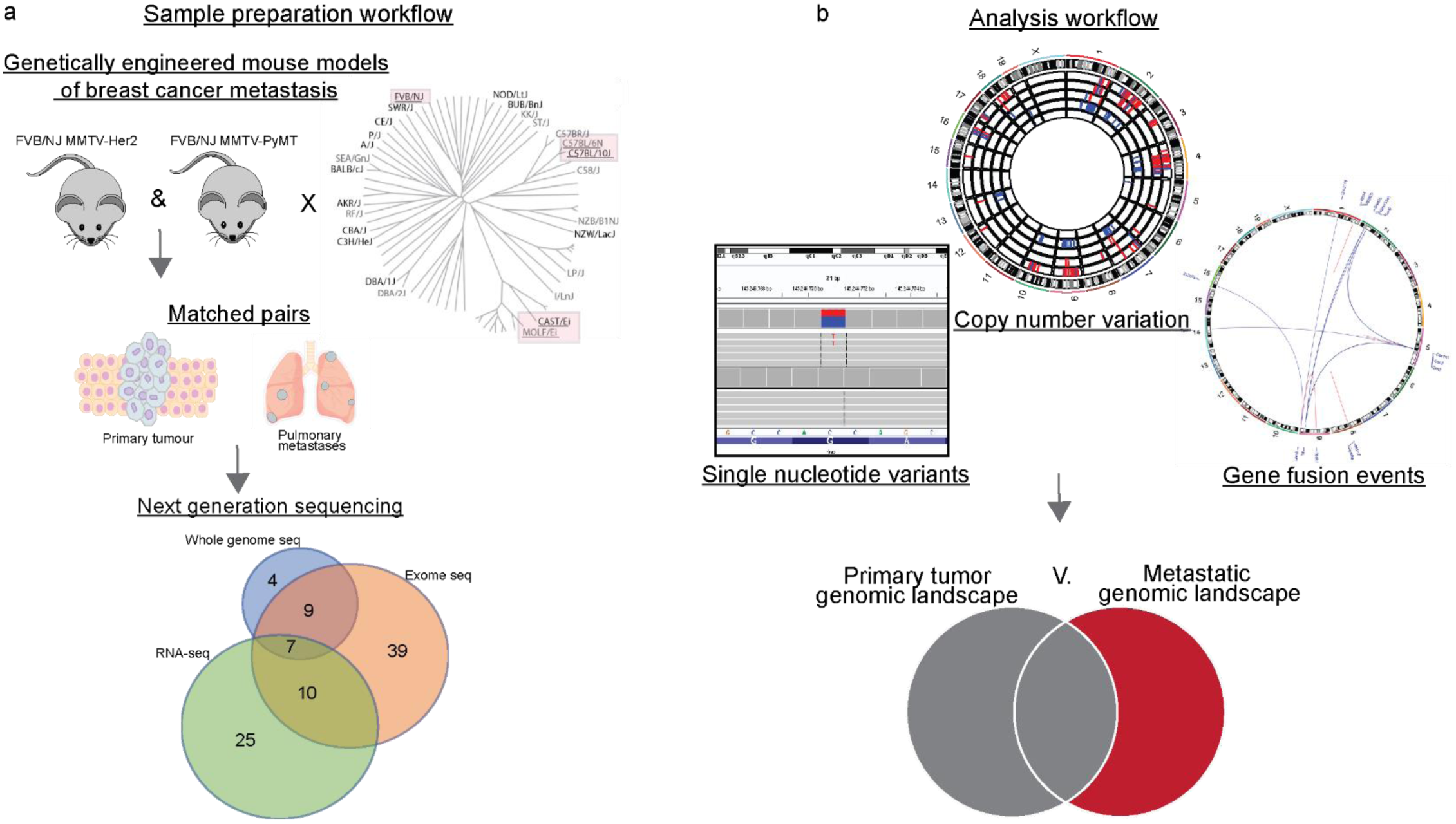
Sample collection and analysis work flow. **a.** Schematic of sample collection for analysis. Preclinical mouse model development followed by sample collection from 14 FVB MMTVxHer2 and 80 out crossed MMTV × PyMT mice. Overlapping analyses performed on paired samples, numbers in Venn diagram represent number of animals. **b.** Schematic of analysis workflow. Next generation sequence data was analysed for single nucleotide variants, copy number variants, and structural variants. The results were then filtered for those events enriched in metastatic tissue only.

**Supplementary Figure 2.**
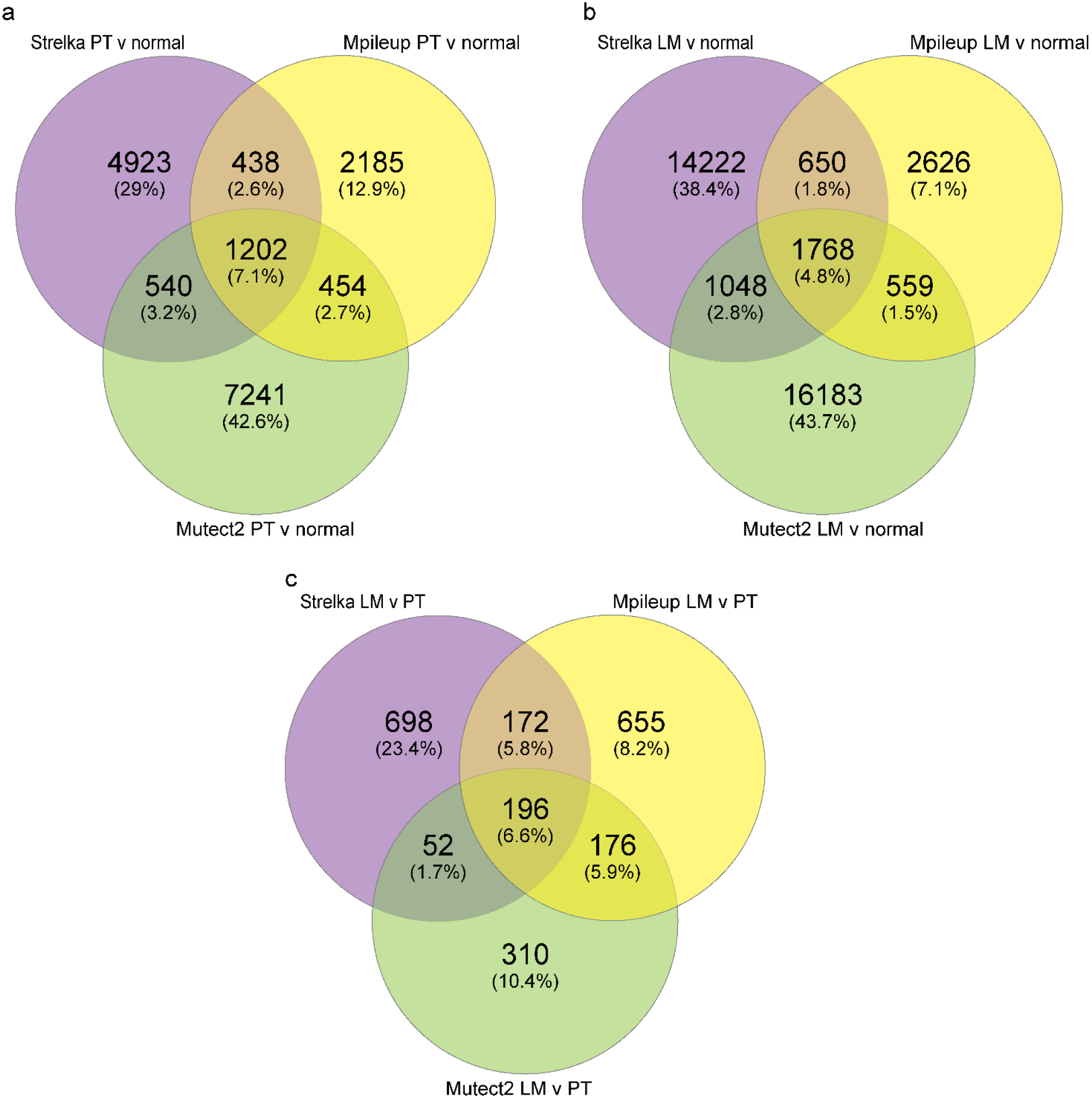
Overlap of three SNV calling algorithms Strelka, Mpileup, and Mutect2 used with exome-seq data collected from primary tumour (PT) and lung metastases (LM) from 65 mice. **a.** SNVs called in primary tumour tissue when compared to normal (strain specific) gDNA. **b.** SNVs called in metastatic tissue when compared to normal (strain specific) gDNA. **c.** SNVs called in metastatic tissue when compared to paired primary tumour tissue using 0.3 allele frequency cutoff.

**Supplementary Figure 3.**
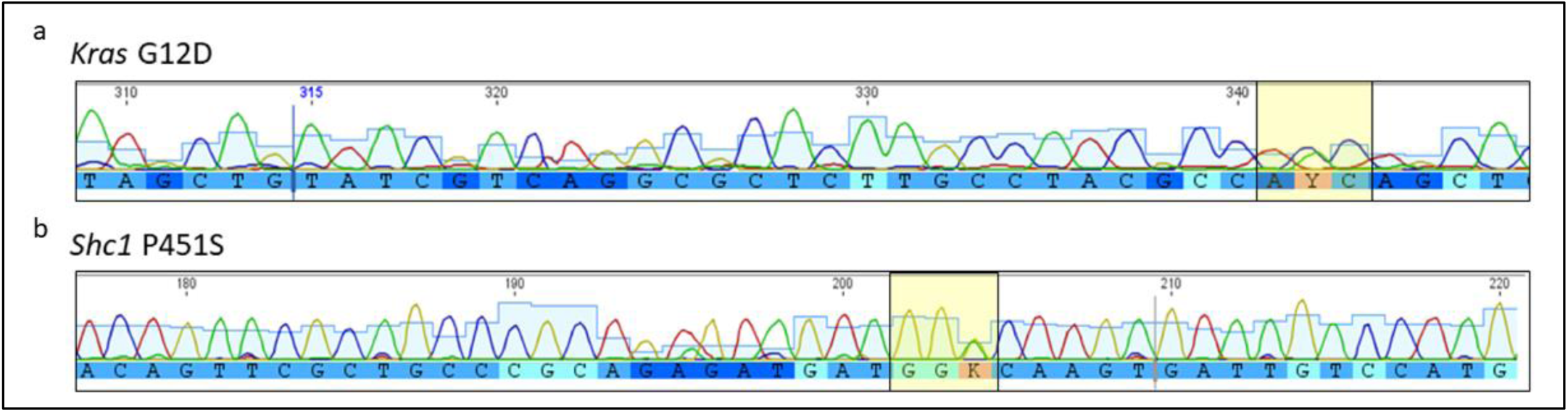
Sanger sequencing spectra showing the validation of SNVs in metastatic gDNA. **a.** C (blue trace)-T (green trace) SNV within the *Kras* gene resulting in the G12D amino acid substitution, Y indicates ambiguity in calling T or C. **b.** G (yellow trace)-T (green trace) substitution within the *Shc1* gene resulting in the P561S amino acid substitution. K indicates ambiguity in calling T or G.

**Supplementary Figure 4.**
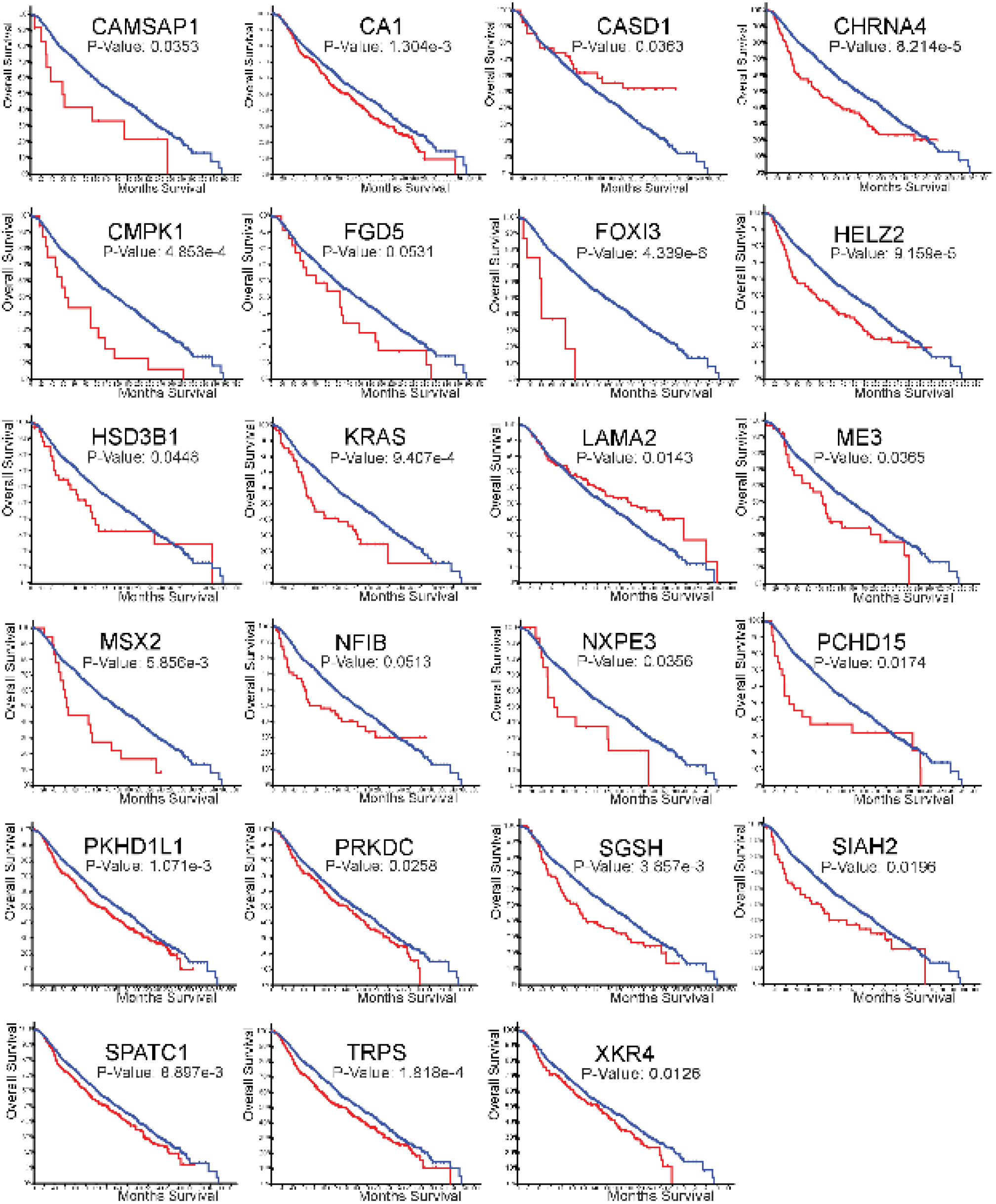
Kaplan-Meier plots generated using METABRIC for 23 genes identified with metastasis-driver SNVs by exome-seq in mice that significantly stratify patient survival when altered in primary tumour tissue (blue = no CNV, red = CNV present).

**Supplementary Figure 5.**
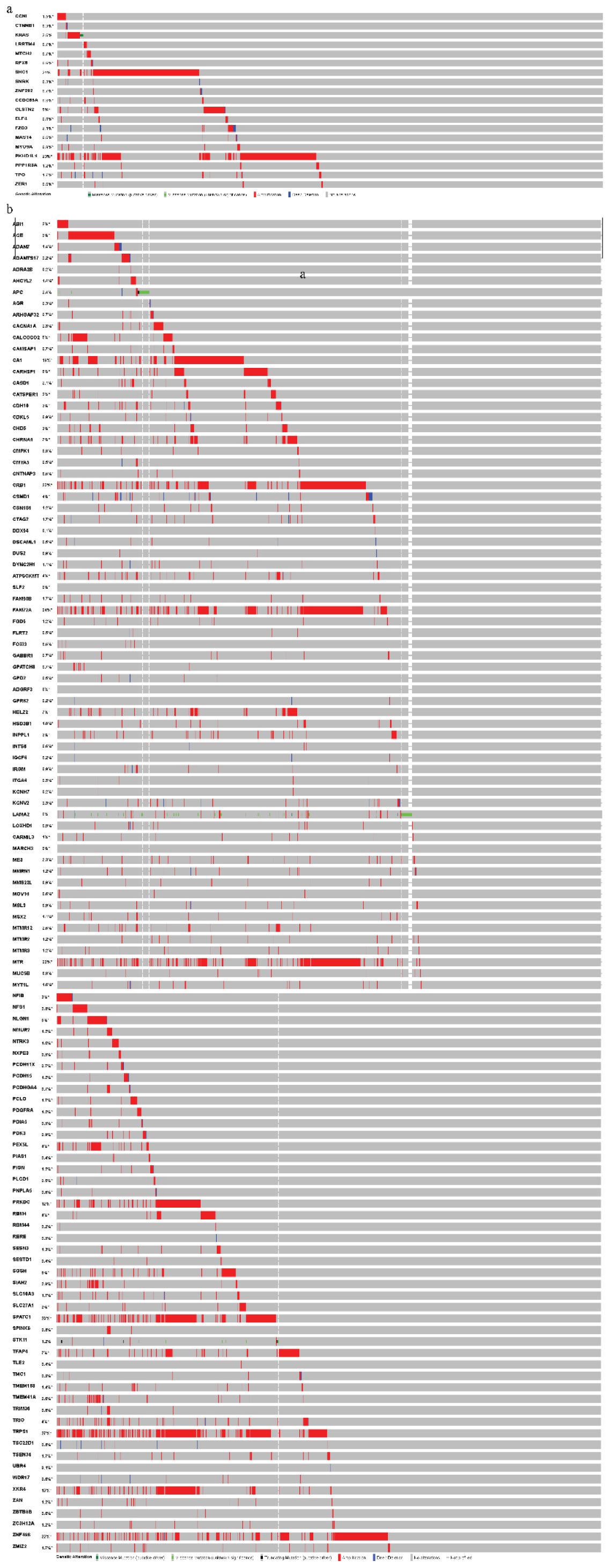
Oncoprint schema from the METABRIC human primary tumour dataset showing copy number variation rates of the **a.** 17 genes with recurrent SNVs and **b.** 147 singly mutated genes identified by exome-seq as putative metastasis-driver mutations (red = amplification, blue = deletion, green = SNV).

**Supplementary Figure 6.**
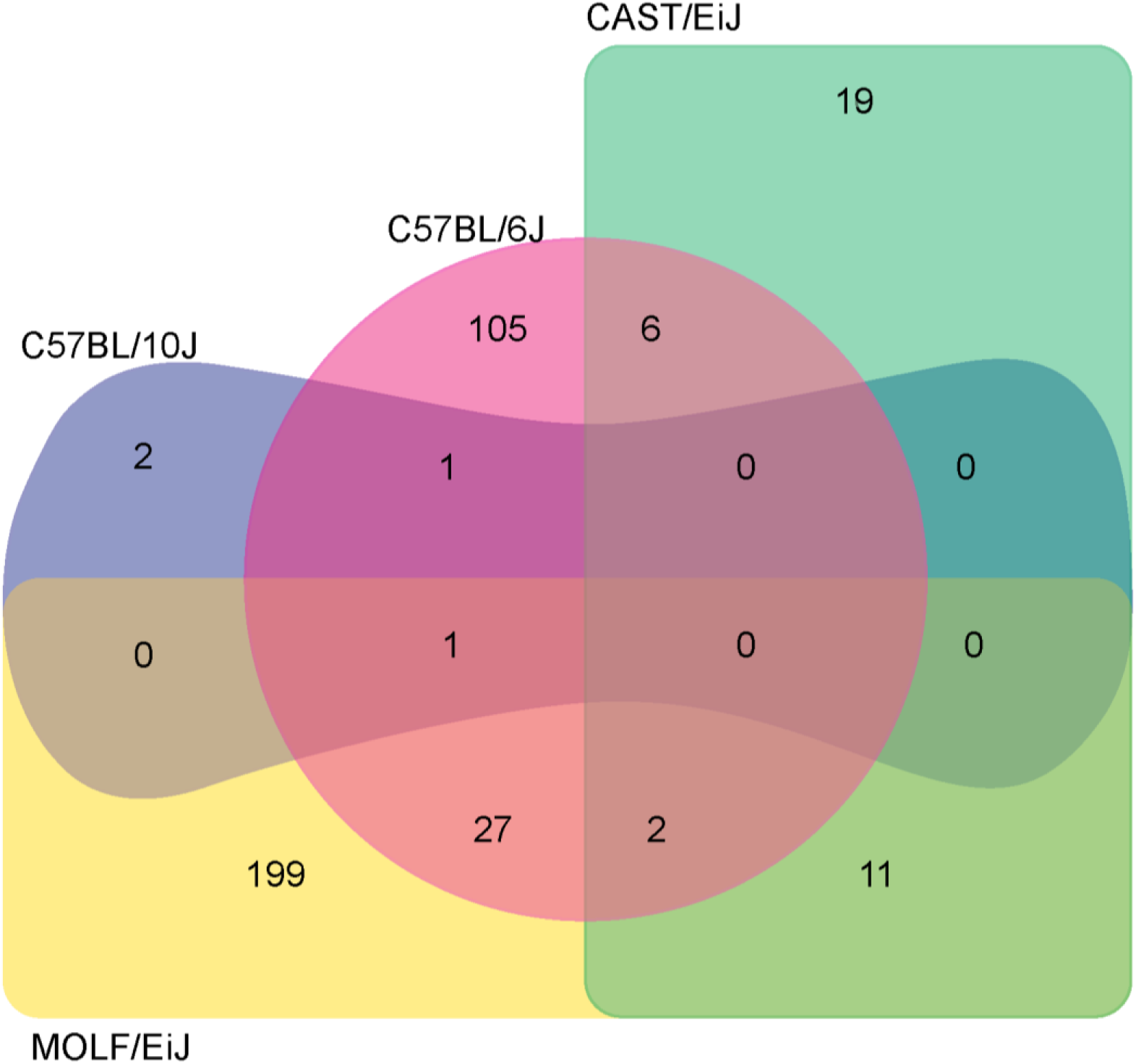
Venn diagram of the genes associated with CNV in each mouse strain. Numbers represent the number of genes, and numbers in overlapping regions represent the number of common CNV-associated CNVs.

**Supplementary Figure 7.**
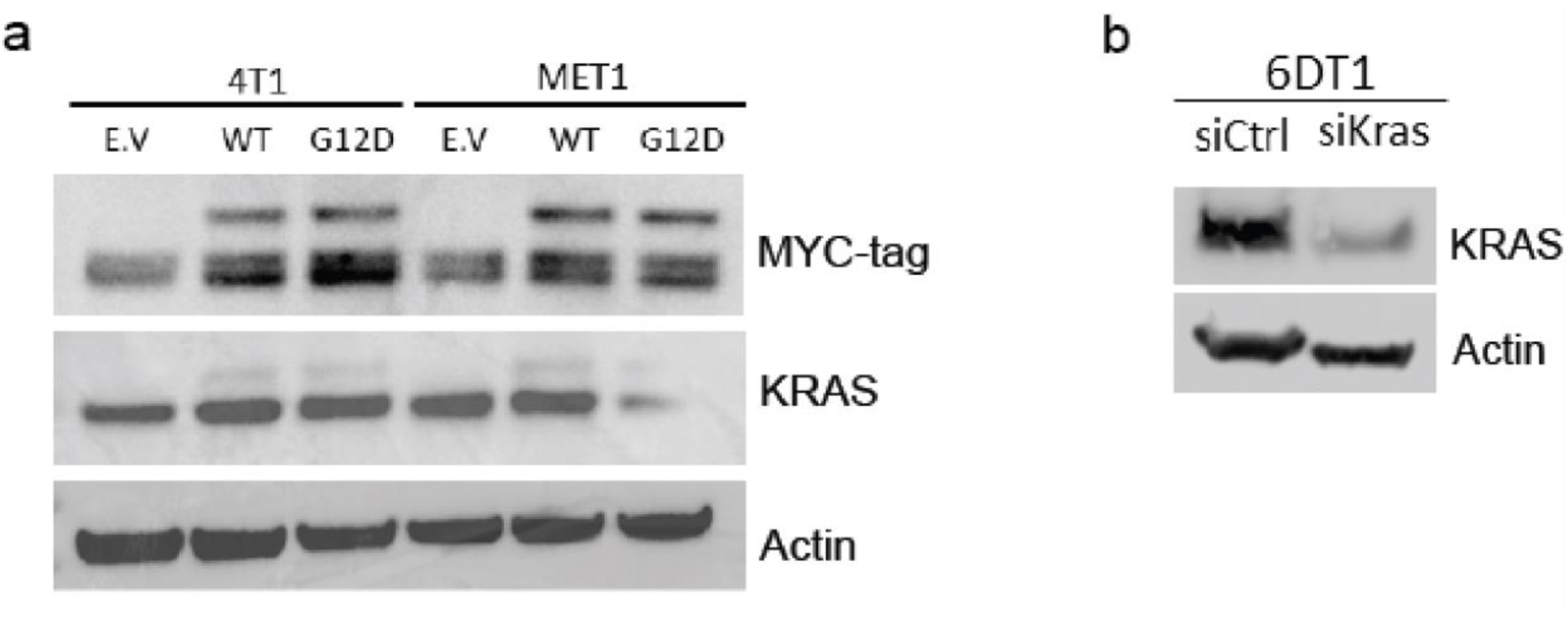
**a.** Western blot showing expression of MYC-tagged KRAS and total KRAS in 4T1 and MET1 transduced with empty vector (EV), *Kras* wildtype (WT), and *Kras* G12D (G12D). **b**. Western blot showing knock down of KRAS in 6DT1 cells 24 hours after transfection siCtrl or si*Kras*.

## Supplementary Tables

**Supplementary Table 1: PyMT and Her2 exome-seq high probability metastasis-specific SNVs**

Sheets: 1. Instances of Her2 metastasis-specific (met. spec.) SNVs, 2. Instances of PyMT met. spec. SNVs, 3. Singly mutated genes, 4. Recurrently mutated genes

Abbreviations: Chr (Chromosome number), Position (mm10 genomic position of mutated SNV), type (mutation type: synonymous, nonsynonymous, or stop gain), alt.fraction (allele fraction within the metastatic tissue), Transcript (NCBI accession number for isoform), Exon (exon harbouring SNV within designated transcript), Codon (codon harbouring SNV within designated transcript), Nuc sub (nucleotide position within designated transcript and substitution), AA sub (amino acid position within gene isoform and resulting substitution)

**Supplementary Table 2: Sequencing validation and overlap**

Sheets: 1. Sanger sequencing (seq.) summary, 2. All seq. summary

Abbreviations: Y (yes), N (no), “/” (and), E (exome seq), R (RNA-seq), W (whole genome seq)

**Supplementary Table 3: PyMT regions of CNV in PT and metastatic tissue compared to normal**

Number of CNV events observed in PT and metastases compared to normal tissue. This table stratifies CNVs by mouse chromosome number and mouse strain. Also listed is the number of animals used in this study per stain, as well as the number of PT or metastatic samples collected from that strain total. Blue cells represent deletion events termed “loss”, and red cells represent amplification events termed “gain”.

**Supplementary Table 4: PyMT regions of CNV and associated genes specific to MOLF/Ei metastatic tissue**

**Supplementary Table 5: PyMT regions of CNV and associated genes specific to CAST/Ei metastatic tissue**

**Supplementary Table 6: PyMT regions of CNV and associated genes specific to C57BL10/nJ metastatic tissue**

**Supplementary Table 7: PyMT regions of CNV and associated genes specific to C57BL6/nJ metastatic tissue**

Sheets:

1. “Strain” loss/gain associated (assoc.) genes, 2. “Strain” loss/gain assoc. pathways

Abbreviations: Name (gene symbol), ID (Term identifier from GREAT ontology), Rank (ordinal rank of the p-value compared to the p-values of other annotations), Raw p-value (uncorrected p-value from the binomial test over genomic regions), FDR q-Value (False discovery rate q-value), Fold Enrichment (fold enrichment of number of genomic regions in the test set with the annotation), Observed Region Hits (actual number of genomic regions in the test set with the annotation), Region Set Coverage (the fraction of all genomic regions in the test set that lie in the regulatory domain of a gene with the annotation,

3. “Strain” recurrent regions of loss or gain

Abbreviations: chr (chromosome), start (position of amplification or deletion start), end (position of amplification or del end), overlap.region (length in bp of overlap in recurrent region of amplification or deletion), freq. (number of individual animals with overlapping region of amplification or deletion), s1 / s2 (region identified in individual animals 1 and 2).

**Supplementary Table 8: PyMT strain summary for CNVs associated genes specific to metastatic tissue** Genes associated with metastasis-specific CNVs and found in more than one mouse strain. Table associated with Supplementary Fig. 5.

**Supplementary Table 9: Fusion events for primary tumour and metastatic tissue compared to normal**

This table shows the data generated from BreakDancer algorithm analysis of 10X WGS from 20 matched pairs of primary and metastatic lesions from FVB/NJ PyMT × MOLF/EiJ and FVB/NJ PyMT x CAST/EiJ animals. We show here the metastasis specific gene fusion data used as input data for Figure 3b. We also she here primary tumour tissue and metastatic tissue gene fusion events versus normal strain specific tissue.

Sheets:

1.Met. spec gene fusion events circos plot input, 2. PT v Norm gene fusion events, 3. Met v Norm gene fusion event

Abbreviations: Sample (mouse ID and tissue type), Chr1 (chromosome where gene 1 is located), Pos1 (position within Chr1 where fusion event break point is located), Chr2 (chromosome where gene 2 is located), Pos2 (position within Chr2 where fusion event break point is located), Type (intra-chromsome events with different strands (ITX) inversion (INV), deletion (DEL), num_Reads (number of reads).

